# Mechanism of non-blocking inhibition of sodium channels revealed by conformation-selective photolabeling

**DOI:** 10.1101/2020.05.05.078071

**Authors:** Mátyás C. Földi, Krisztina Pesti, Katalin Zboray, Tamás Hegedűs, András Málnási-Csizmadia, Peter Lukács, Arpad Mike

## Abstract

Sodium channel inhibitor drugs can exert their effect by either blocking, or modulating the channel. The extent of modulation versus channel block is crucial regarding the therapeutic potential of drug candidates. Modulation can be selective for pathological hyperactivity, while channel block affects vital physiological function as much as pathological activity. Previous results indicated that riluzole, a drug with neuroprotective and antiepileptic effects, may have a unique mechanism of action, where modulation is predominant, and channel block is negligible. We studied the effects of riluzole on rNa_V_1.4 channels expressed in HEK cells. We observed that inhibition by riluzole disappeared and reappeared at a rate that could not be explained by association/dissociation dynamics. In order to verify the mechanism of non-blocking modulation, we synchronized photolabeling with the voltage clamp protocol of patch-clamp experiments. Using this method, we could bind a photoreactive riluzole analog covalently to specific conformations of the channel. Photolabeling was ineffective at resting conformation, but effective at inactivated conformation, as judged from persisting modulated gating after removal of unbound photoactive drug from the solution. Mutation of the key residue of the local anesthetic binding site (F1579A) did not fully prevent ligand binding and inhibition, however, it eliminated most of the modulation caused by ligand binding. Our results indicate that riluzole binds with highest affinity to the local anesthetic binding site, which transmits inhibition by the unique non-blocking modulation mechanism. Our results also suggest the existence of one or more additional binding sites, with lower affinity, and different inhibition mechanism.

## Introduction

### Hyperexcitability and state-dependent drugs

Hyperexcitability is at the core of a rather diverse set of disorders affecting the heart, skeletal muscles and the nervous system. Out-of-control electric activity is involved in the development of several types of epilepsies, chronic pain syndromes, neuromuscular disorders, cardiac arrhythmias, and even psychiatric disorders^1^. These conditions can originate from genetic conditions (mutations altering the operation of channels themselves, or proteins involved in their modulation), or can be due to damages caused by mechanical injury, inflammation or ischemia.

To suppress hyperexcitability, sodium channels are the primary target, because they are responsible for the fast onset of action potentials, as well as their all-or-none and self-regenerating nature.

In order to design therapeutically useful sodium channel inhibitors, however, one encounters the seemingly impossible task of having to prevent pathological hyperexcitability, while maintaining normal physiological activity of nerves and muscles. Interestingly, there are compounds, which are able to carry out this feat – at least to a certain extent. These compounds include antiarrhythmic, antiepileptic, antispastic etc. compounds. The trick that enables them to do so is state-dependence, i.e., their preference for certain conformational states of the channel protein. Most sodium channel inhibitors prefer inactivated state to resting state, they bind to it more rapidly, and/or dissociate from it more slowly. The fact that an inhibitor has higher affinity to inactivated state, means that drug-bound inactivated channels form an energetically favorable complex. On the one hand this means slower dissociation of the ligand from this conformation, and on the other hand it also inevitably means that inactivated conformation of the channel is stabilized by drug binding, therefore recovery from inactivated to resting state will be slower. This effect – called modulation of channel gating – is an inseparable element of state-dependent inhibition, as described by the modulated receptor hypothesis^2^. High affinity to inactivated state ensures delayed dissociation, while modulation ensures delayed conformational transition to the low affinity resting state, thus restraining both possible pathways to recovery (dissociation followed by recovery from inactivated state, or recovery from inactivated state followed by dissociation).

Pathological states induced by injury, inflammation, ischemia, tumor, or epilepsy alter the electrical characteristics of excitable cells, which may include a depolarized membrane potential due to energy failure, increased leakage currents, left-shifted voltage sensitivity of sodium channels, and increased persistent component of the sodium current^3–8^. These changes make sodium channels more likely to be in open and inactivated conformations, therefore state-selectivity alone is enough for preferential inhibition of pathological tissue. On a first impression one could suppose that the stronger the state-preference is, the better the drug will be.

However, the temporal aspect must also be considered. Action potentials are fired repetitively, and pathological behavior of neurons is often manifested as high frequency firing. The extent of state-dependence is, therefore, not the only crucial aspect, equally important is the onset/offset dynamics of state dependent binding. Its significance is obvious in the case of Class 1 antiarrhythmics, where subclasses a, b, and c differ in their association/dissociation kinetics, but the same is true for the much higher firing rates in central and peripheral neurons. For selective inhibition of cells firing at pathologically high frequency, the ideal drug should work as a low pass filter, with a steep frequency response, to be able to distinguish pathological and physiological rates of firing. This, however presents a theoretical limit for state dependent binding, because fast dissociation precludes high affinity. One would want both fast binding/unbinding dynamics, and high state dependence. However, high state-dependence requires high affinity to inactivated state, and high affinity means slow dissociation, which means that binding/unbinding dynamics cannot be fast.

Intriguingly, riluzole seems to be able to elude this limitation. In this study we examined how this is possible.

We first describe the peculiar pattern of inhibition we observed during and after riluzole perfusion. We designed a voltage-clamp protocol where block and modulation of the channels are monitored in parallel. Paradoxically, two distinct recovery processes coexisted, with rates differing by more than two orders of magnitude. We presumed that this peculiar feature may be the key to being able to feature fast kinetics and high affinity at the same time. A fast recovery allowed channels to regain their ability to conduct ions within ~10 ms. In spite of their ability to conduct, however, channels remained modulated by the drug for a much longer time: it took ~2 s for the channels to recover from the modulatory influence. This mechanism enables riluzole to function as a low pass filter with an exceptionally steep frequency response, which is probably a key element of its distinctive therapeutic efficacy.

We aimed to identify the physical processes that underlie fast and slow recovery processes.

## Results

### The peculiar effect of riluzole – two distinct recovery processes

#### Recovery in the presence of the drug

Investigating the “frequency response” of riluzole in our previous studies^9,10^ we found it to be exceptionally good, in fact it was so steep, that it exceeded the theoretically possible limit (this rate of dissociation, with the affinity of the compound would require impossibly fast association; see Discussion). Of the >70 sodium channel inhibitor drugs we have investigated thus far (unpublished data), no other had such steep frequency response. Conventionally in voltage clamp experiments frequency characteristics of inhibition are assessed using the “recovery from inactivation” (**RFI**) protocol, which measures the rate of recovery at a constant hyperpolarized membrane potential.

As illustrated in Fig. 1, this consists of pairs of depolarizations, where the length of the hyperpolarizing gap separating them is progressively increased. In general, sodium channel inhibitor drugs are most effective at short hyperpolarizing gaps (which occurs at high frequency firing), and their inhibitory effect gradually decreases with longer hyperpolarizations. This is conventionally explained by the progressive dissociation of drug molecules from resting state throughout the hyperpolarizing period, because they have lower affinity to resting than to inactivated state. In the absence of any drug, at −130 mV channels recovered with a time constant of 0.39 ± 0.08 ms (exponentials on the 3^rd^ power, see Methods). In the presence of 100 μM riluzole recovery was delayed, it proceeded with a time constant of 2.25 ± 0.004 ms (p = 0.00012, paired t-test, n = 9). Does this time constant reflect dissociation?

**Fig. 1.**
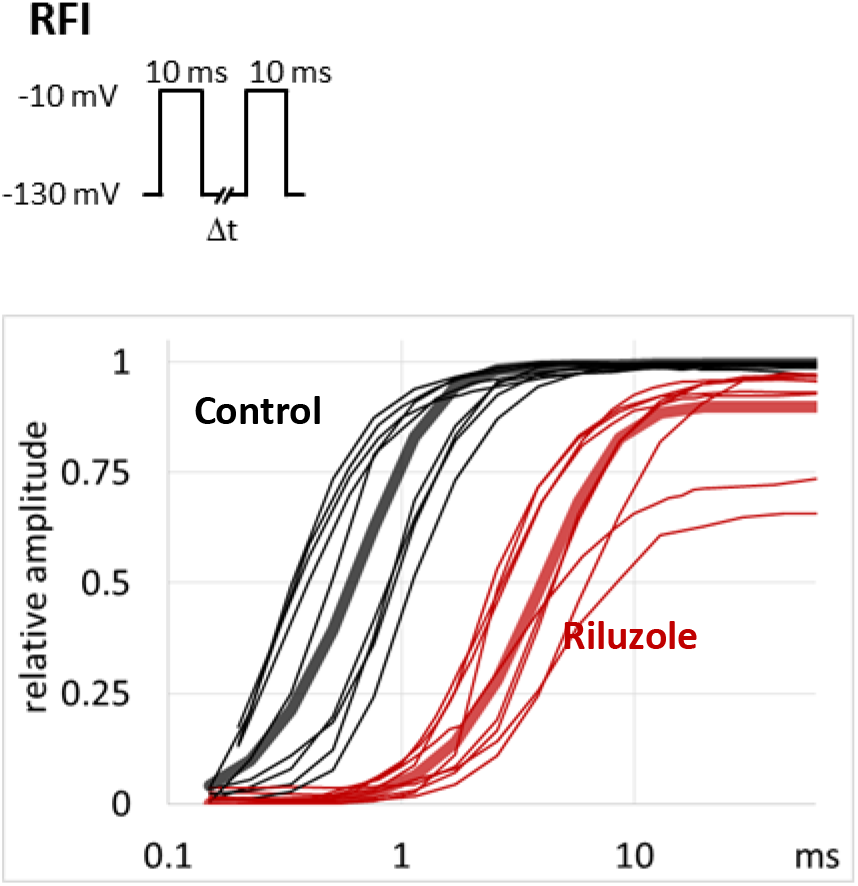
Recovery from inactivation in the absence and presensce of 100 μM riluzole in WT channels. The RFI protocol is shown in the upper panel. Duration of the hyperpolarizing gap between the two depolarizations is progressively increased. Peak amplitudes of the 2nd pulse-evoked currents are plotted against hyperpolarizing gap duration. Thin black lines – normalized control amplitudes in n = 9 cells), thin red lines – amplitudes in the same 9 cells in the presence of 100 μM riluzole. Data from each cell was normalized to its own control. Averaging (thick lines) was done as described in Methods.

#### Recovery upon removal of the drug

We investigated onset and offset kinetics with a protocol, which consisted of 100 successive trains of three depolarizations; we will call it “three pulse train” (**3PT**) protocol (Fig. 2a and d).

**Fig. 2.**
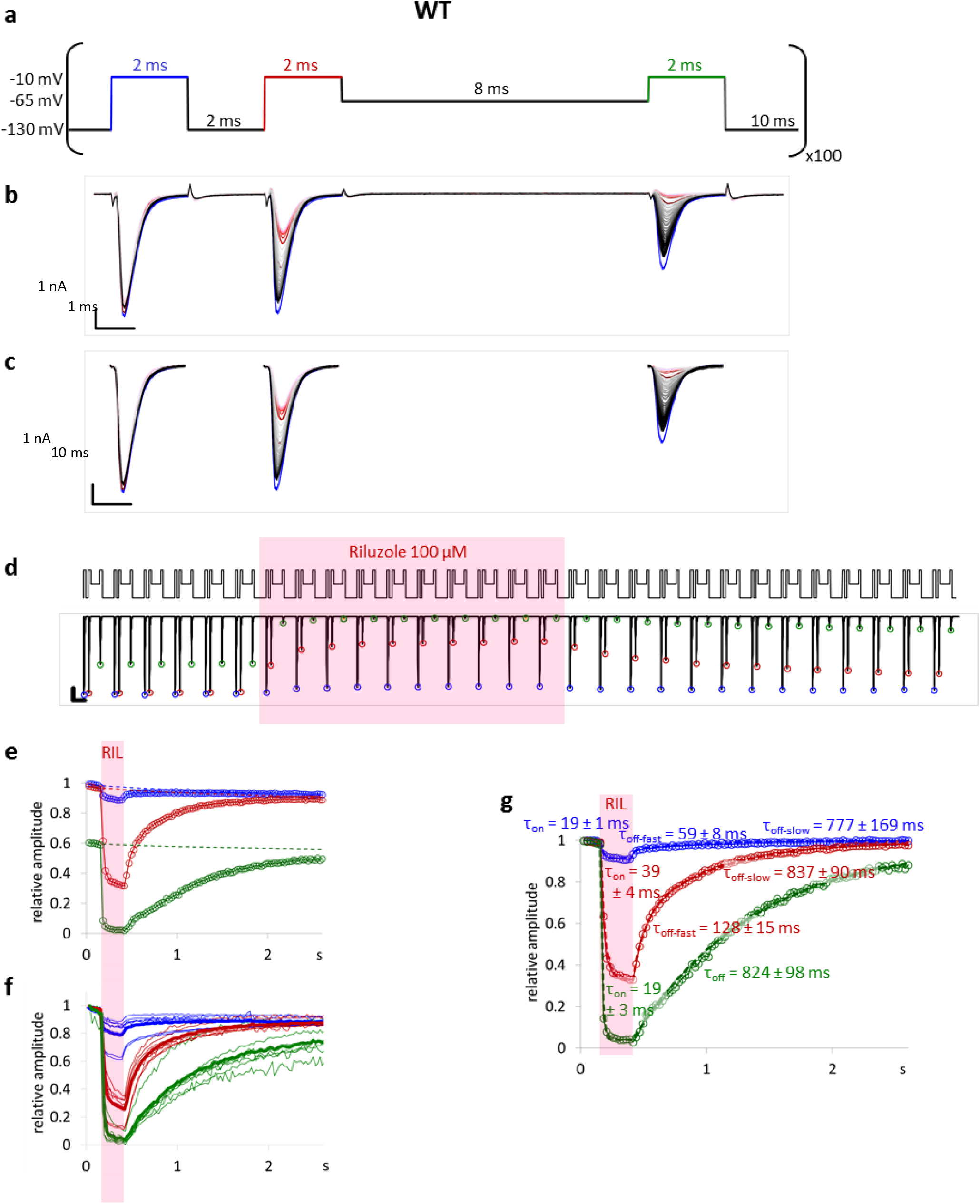
Effect of riluzole perfusion as monitored with the 3PT protocol on WT channels. **a** The voltage protocol. Blue, red and green color indicates 1^st^, 2^nd^, and 3^rd^ depolarizations, respectively, as in panels **d** to **g**. **b** Example for sodium currents evoked in a typical WT channel expressing cell by the three depolarizations of the 3PT protocol. Subsequent traces are overlaid on each other. Control traces are shown in blue, traces during riluzole application are shown in dark to light red, traces recorded during washout are shown in light gray to black. **c** The same currents after subtraction of capacitive and leakage artifacts. **d** Illustration of the first 29 consecutive trains (protocol in the upper panel), and the currents evoked by them (lower panel). Peaks of 1^st^, 2^nd^, and 3^rd^ pulse-evoked currents are marked with blue, red, and green circles, respectively. Shaded area indicates the perfusion of 100 μM riluzole. **e** Plot of peak amplitudes throughout the whole experiment, normalized to the amplitude of the first evoked current. Each circle represents the peak amplitude of a current evoked by either 1^st^ (blue), 2^nd^ (red), or 3^rd^ (green) pulses of each consecutive train. Dashed lines show the progression of slow inactivation during the experiment. **f** Amplitude plots for 7 individual cells. To help comparing the extent of inhibition, 1^st^, 2^nd^, and 3^rd^ pulse-evoked currents were normalized each to its own control (peak amplitudes recorded during the first train). **g** Onset and offset time constants (mean ± SEM) for the three different peak amplitude plots. Mono- or bi-exponential functions were fitted to traces after correcting for slow inactivation. Dashed lines show exponentials fitted to this particular cell. Time constant values show mean ± SEM of time constants fitted for the 7 individual cells shown in panel **f**.

This protocol was similar to the one we called by this name before^9^; but in this study it was optimized for maximal time resolution. The protocol is designed to monitor changes in both gating kinetics and gating equilibrium. For the sake of high time resolution, it uses only a single interpulse interval: 2 ms (instead of several interpulse intervals as in **RFI)**, and only a single membrane potential: −65 mV (instead of several membrane potentials as in **SSI**). In contrast to the previously used 333 ms resolution (3 Hz), the protocol we currently used could record changes at 26 ms resolution (38.5 Hz). In each experiment we delivered 100 consecutive trains: after 6 initial control trains we perfused riluzole throughout 10 trains (i.e., for 260 ms), and then monitored washout for the remaining 84 trains. Perfusion was started and stopped during the 10 ms intervals between two trains, which was amply enough for complete exchange of the solution with the theta tube system (see Methods).

Recordings of all 100 trains from a typical cell are shown in Fig. 2b. The 6 control traces are printed in blue, the 10 traces recorded during riluzole perfusion are in red (dark to light red indicates consecutive traces), and currents evoked during washout are shown in light gray to black. Capacitive and leakage artifacts were subtracted (Fig. 2c), and the corrected peak amplitudes were determined for each pulse. In Fig. 2d the same corrected current traces are shown in sequential order, together with the voltage protocol, as they were evoked in the experiment (only the first 29 trains, for the sake of clarity). Peak amplitudes are marked by circles: blue (1^st^ pulses), red (2^nd^ pulses) and green (3^rd^ pulses). In Fig. 2e, 1^st^, 2^nd^ and 3^rd^ pulse-evoked peak amplitudes were plotted separately against time for the same cell as shown in Fig2b to d. All amplitudes were normalized to the 1^st^ pulse-evoked amplitude at the start of the experiment. In Fig. 2f, 1^st^, 2^nd^, and 3^rd^ pulse-evoked amplitudes for n = 7 cells are shown; amplitudes were normalized each to its own control, in order to help comparison of inhibited fractions. Current traces for the same 7 cells are shown in Supplementary Fig. 1, together with a detailed discussion of fitted time constants.

A small fraction of channels (11.4 ± 0.1 % by the end of the 100^th^ train) underwent slow inactivation during the test; this was a necessary trade-off for high temporal resolution. To correctly calculate the extent of inhibition, the effect of slow inactivation (shown as dashed lines in Fig. 2e) had to be subtracted (as described in Methods). This is illustrated in Fig. 2g, which is the exact same data as in Fig. 2e, after correction. Measurement of the extent of inhibition, and exponential fitting of onset and offset were done on the corrected plots.

The offset of inhibition upon actual removal of the drug was more than hundredfold slower than recovery from inactivation in the presence of riluzole: it occurred with a time constant of 824 ± 99 ms (Fig. 2g). Paradoxically, within each cycle the inhibition seemed to have disappeared during each 10 ms of between-trains hyperpolarization (see blue traces), and then was re-established before the 2^nd^ depolarization, and even more during the subsequent 8 ms spent near the V_1/2_ (−65 mV), causing a massive inhibition at the 3rd pulse of each train. This pattern repeated itself during riluzole perfusion, and also for 1-2 s after riluzole had been washed out.

The extent of inhibition in 1^st^, 2^nd^, and 3^rd^ pulse-evoked currents the inhibition was 16.1 ± 4.0 %, 73.4 ± 3.9 %, and 95.4 ± 1.2 %. Apparent affinity (IC_50_) values can be determined from the extent of inhibition, as described in Methods; these values corresponded with 606 ± 137 μM, 33.5 ± 6.8 μM, and 3.98 ± 1.32 μM, respectively.

It is evident from Fig. 2 that two distinct recovery processes coexist, with completely different kinetics. The fast recovery process was repeatedly completed between trains of the **3PT** protocol (i.e., within 10 ms), even though riluzole was being continuously perfused. However, after washing out riluzole, the recovery from riluzole’s effect took a few seconds, as seen in Fig. 2g. Recovery is conventionally explained by dissociation, but the two recovery processes with more than hundredfold different time constants obviously cannot both reflect dissociation. To investigate what physical processes underlie the progression of both fast and slow recovery, we first studied the effect of mutating the most important residue of the local anesthetic binding site, F1579. The presence of this aromatic residue has been shown to be crucial in determining the affinity of binding^11,12^, and also in coupling drug binding to altered voltage sensor movements^13,14^, which is the basis of gating modulation.

#### The two recovery processes in binding site mutant channels

We repeated the same experiments, using the **RFI** and **3PT** protocols in cells expressing binding site mutant (F1579A) channels. We found that both the fast and the slow processes were radically altered by the mutation. In mutant channels fitting the recovery required a biexponential function, the major fast component had a time constant of 0.30 ± 0.02 ms, and it contributed 85.5 ± 2.5 % to the amplitude. Time constant of the minor, slow component was 4.36 ± 0.62 ms. Riluzole only slightly delayed fast component, to 0.43 ± 0.02 ms (p = 0.012, paired t-test, n = 7) (Fig. 3). The minor, slow component was not changed significantly.

**Fig. 3.**
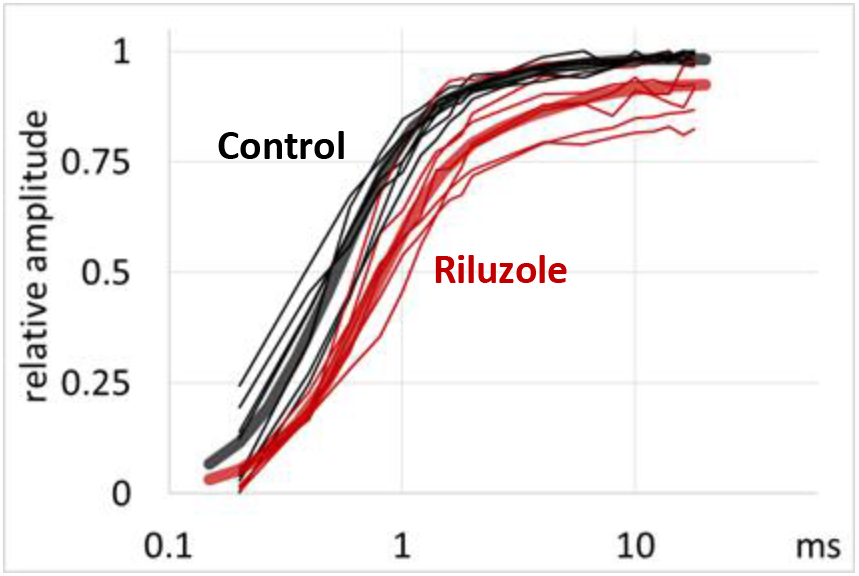
Recovery from inactivation in the absence and presensce of 100 μM riluzole in F1579A mutant channels. Peak amplitudes are plotted against hyperpolarizing gap length. Thin black lines – normalized control data from n = 7 cells, thin red lines – 100 μM riluzole from the same 7 cells, normalized each to its own control. Averaging (thick lines) was done as described in Methods.

The effect of mutation on the fast recovery was also indicated by the decreased inhibition of 2^nd^ pulse-evoked currents in the **3PT** protocol (Fig. 4), see the minimal difference between blue and red traces in Fig. 4f. Interestingly, 3^rd^ pulse-evoked currents were still considerably inhibited (by 37.2 ± 6.6 %). Apparent affinity (IC_50_) values, calculated from the extent of inhibition, were 1590 ± 180 μM, 895 ± 126 μM, and 114 ± 20.0 μM for 1^st^, 2^nd^, and 3^rd^ pulse-evoked currents, respectively. Compared to the apparent affinity values of WT channels, 1^st^ pulse-evoked responses were least sensitive to the mutation (2.62-fold decrease in affinity), while 2^nd^ and 3^rd^ pulse-evoked responses much more sensitive (26.7-fold and 28.6-fold decrease in affinity, respectively).

**Fig. 4.**
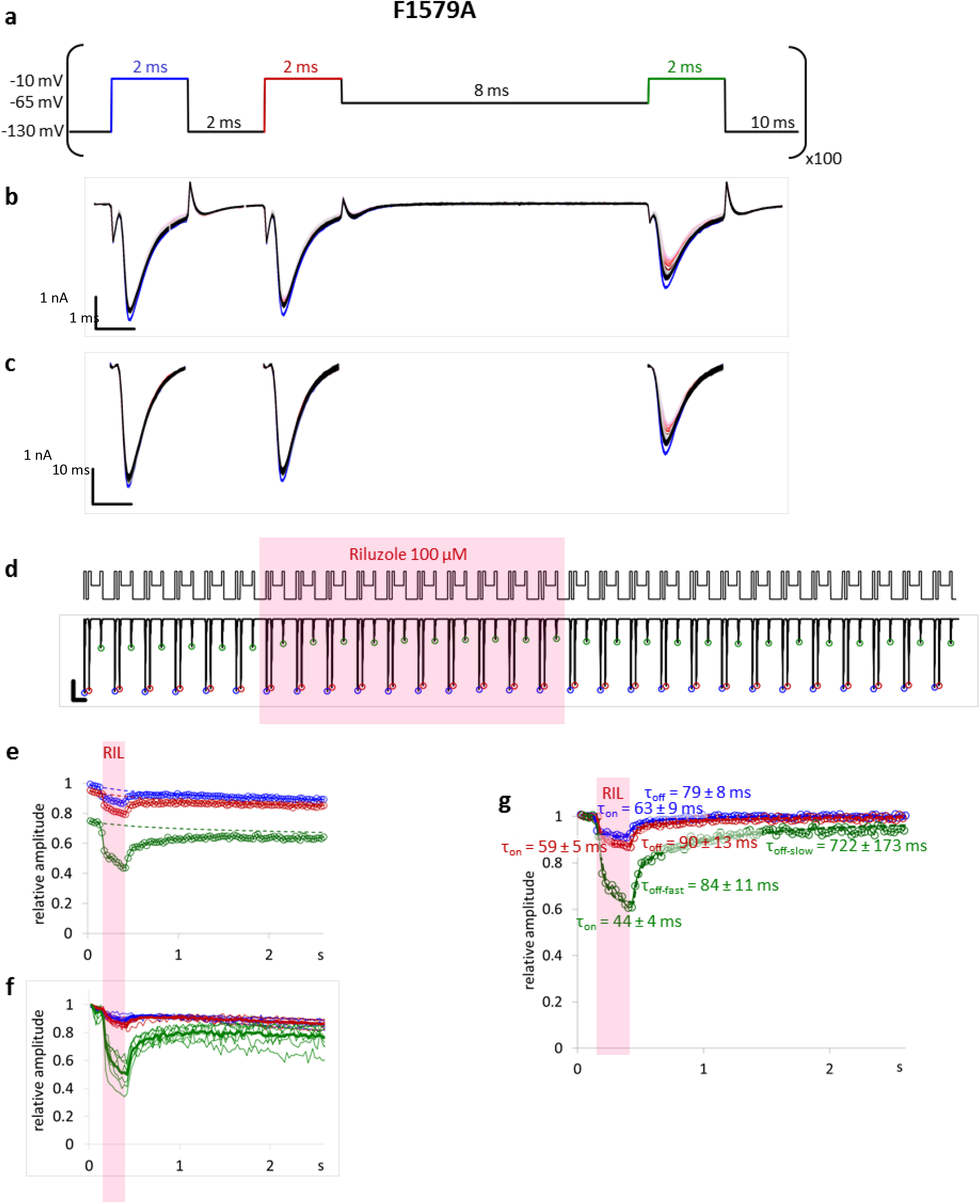
Effect of riluzole perfusion as monitored with the 3PT protocol on F1579A mutant channels. **a** The voltage protocol. Blue, red and green color indicates 1^st^, 2^nd^, and 3^rd^ depolarizations, respectively, as in panels **d** to **g**. **b** Example for sodium currents evoked in a typical F1579A mutant channel expressing cell by the three depolarizations of the 3PT protocol. Subsequent traces are overlaid on each other. Control traces are shown in blue, traces during riluzole application are shown in dark to light red, traces recorded during washout are shown in light gray to black. Scale bar: 1 nA, 1 ms. **c** The same currents after subtraction of capacitive and leakage artifacts. **d** Illustration of the first 29 consecutive trains (protocol in the upper panel), and the currents evoked by them (lower panel). Peaks of 1^st^, 2^nd^, and 3^rd^ pulse-evoked currents are marked with blue, red, and green circles, respectively. Shaded area indicates the perfusion of 100 μM riluzole. Scale bar: 1 nA, 10 ms. **e** Plot of peak amplitudes throughout the whole experiment, normalized to the amplitude of the first evoked current. Each circle represents the peak amplitude of a current evoked by either 1^st^ (blue), 2^nd^ (red), or 3^rd^ (green) pulses of each consecutive train. Dashed lines show the progression of slow inactivation during the experiment. **f** Amplitude plots for 9 individual cells. To help comparison between 1^st^, 2^nd^, and 3^rd^ pulse-evoked currents, as well as between WT (Fig. 2) and mutant channels, 1^st^, 2^nd^, and 3^rd^ pulse-evoked currents were normalized each to its own control (peak amplitudes recorded during the first train). **g** Onset and offset time constants (mean ± SEM) for the three different peak amplitude plots. Mono- or bi-exponential functions were fitted to traces after correcting for slow inactivation. Dashed lines show exponential functions fitted to this particular cell. Time constant values show mean ± SEM of time constants fitted for the 9 individual cells shown in panel **f**.

The slow recovery process was also accelerated, (compare Fig. 4e,f,g to Fig. 2e,f,g). The major time constant decreased approximately tenfold, from 823 ms to 128 ms. For a detailed analysis of time constants see Supplementary Fig. 1.

### Conformation-selective photolabeling

#### The problem of interrelated binding and gating

To summarize, thus far we have learned that there were two distinct recovery processes, and they were both accelerated by mutation of the local anesthetic binding site. What physical processes may underlie them? In the presence of riluzole the rate of the fast recovery process may be limited either by modulated gating (i.e., the recovery from inactivation is slowed down by the bound drug), or by the dissociation of the drug – which is the conventional explanation.

The modulated receptor hypothesis^2^ predicts that, since the drug has different affinities to different conformational states, drug binding must alter the energetics of conformational transitions, making higher affinity states more stable. Any mutation that changes binding affinities to different states must also change the effect of the drug on rates of transitions between these states (i.e., the gating of drug-bound channels). In other words, decreased affinity must necessarily cause decreased modulation of gating. Mutation-induced changes in the fast and slow recovery processes, therefore, could both be manifestations of decreased affinity, on the level of gating and binding, respectively. Mutation-induced acceleration of the fast recovery process (compare Fig. 1 and Fig 3) may be due to decreased modulation of channel gating, while acceleration of the slow recovery process (compare Fig. 2f and 4f) may be caused (at least partly) by faster ligand unbinding. If we accept this explanation, we must suppose that channel gating alone (without dissociation of the inhibitor) can make the channel available for activation and conduction, in other words “non-blocking modulation"^9^ is possible.

The complex problem of interrelated gating and binding/unbinding can be simplified using photolabeling-coupled electrophysiology, as we have previously demonstrated^9^. By binding the photoreactive riluzole analog, azido-riluzole, covalently to the channel, we could exclude the processes of binding and unbinding. We found that covalently bound channels were still able to conduct ions, but with modulated gating. The extent of modulation (delay in recovery from inactivation) almost exactly matched the one induced by the presence of riluzole (see Fig. 4e in ref^9^).

We have not investigated, however, whether this was based on specific binding to the local anesthetic binding site, and whether binding was conformation-dependent. In this study we have improved the method of photolabeling-coupled electrophysiology by synchronizing UV light pulses with the voltage protocol, instead of the continuous illumination we had used previously^9^. This allowed us to target specific conformations of the channel, and also to investigate if gating transition to the unfavorable resting conformation was immediately followed by unbinding. This way these data could help us to identify the physical processes underlying both fast and slow recovery processes.

#### State-dependence of binding

Riluzole, like most sodium channel inhibitors, is known to preferentially bind to inactivated channels^15^. We first intended to investigate how the conformation of the channel influences the ability of the drug to bind. Gating kinetics and equilibrium of the channels was investigated both before drug perfusion and after washout, using the “recovery from inactivation” (**RFI**), “steady-state inactivation” (**SSI**) and “state-dependent onset” (**SDO**) voltage protocols, as we have described earlier^9^. During drug perfusion we used one of three illumination protocols, in order to compare the possibility of binding to different conformations (Fig. 5a). All three protocols were repeated at 2.5 Hz (every 400 ms), and within each repetition a 90 ms UV light pulse was applied. The UV pulses were delivered either during hyperpolarizations, ("resting-state-illumination"); during depolarizations ("inactivated-state-illumination"); or while cells were kept at the approximate half-inactivation voltage ("V-half-illumination"), i.e., when channels were distributed roughly equally between the two conformations. To each cell, a maximum of 450 UV pulses were delivered (*i.e.*, for 3 min), it was stopped earlier whenever evoked currents were inhibited to ~20% of their original amplitudes. (It is important to note that an 80% inhibition did not mean an 80% occupancy of binding sites, as we will discuss below.)

**Fig. 5.**
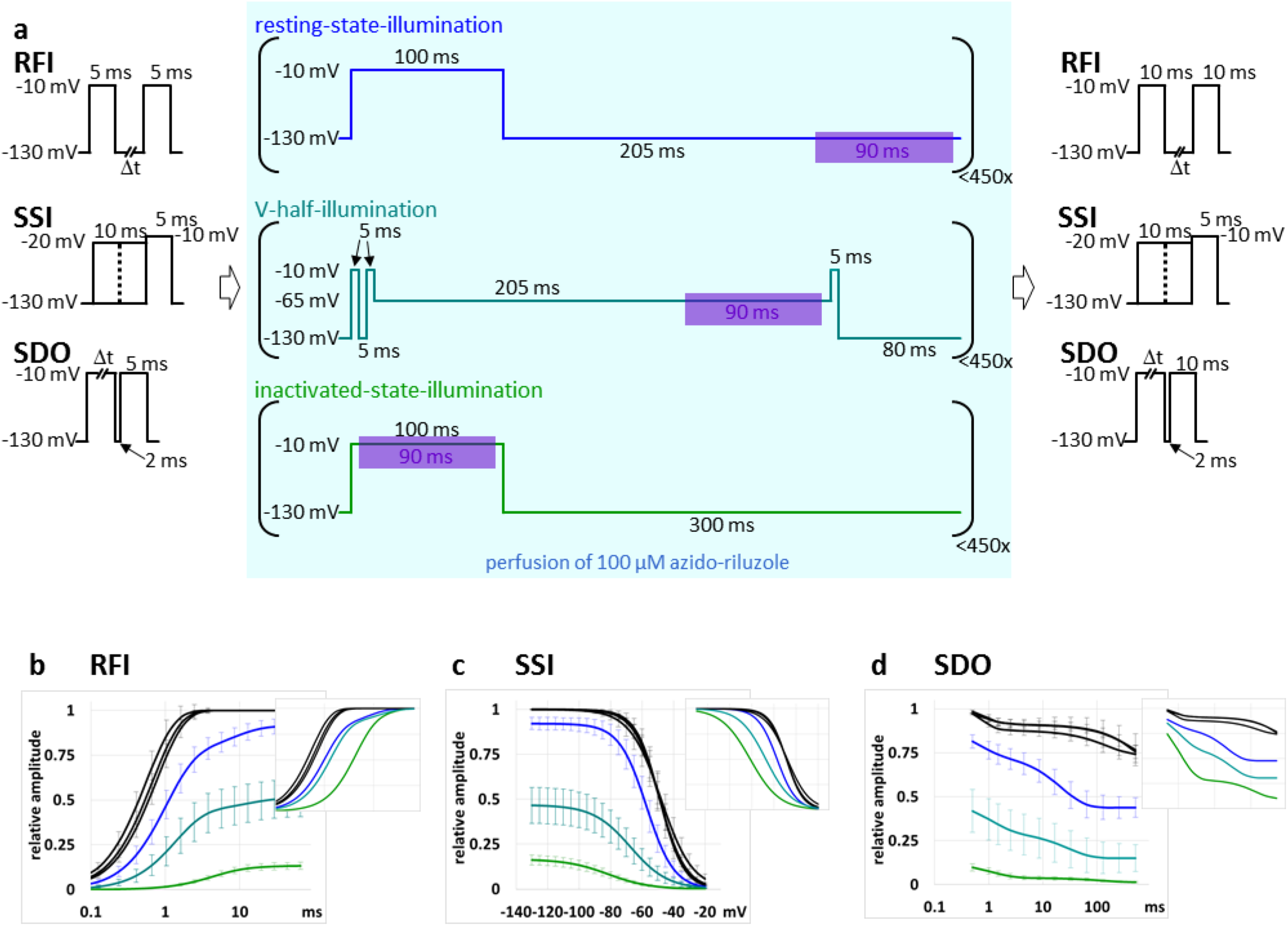
Effect of conformation-selective photolabeling by azido-riluzole on gating kinetics of WT channels. **a** Experimental protocol. **RFI**, **SSI**, and **SDO** protocols were run before and after azido-riluzole perfusion and pulsed UV illumination. During azido-riluzole perfusion 90 ms UV illumination pulses were used (shown by purple shaded areas) in each of the three voltage protocols. **b, c, d** Assessment of gating kinetics and equilibrium before and after azido-riluzole perfusion and UV irradiation. Black lines show control data before azido-riluzole perfusion and UV illumination, colored lines show data measured after stopping UV pulses and washing out azido-riluzole. Blue lines: after resting-state-illumination protocol; teal lines: after V-half-illumination protocol; green lines: after inactivated-state-illumination protocol. Amplitudes were normalized to the maximal amplitude in control. Insets: Amplitudes normalized each to its own maxima. **d:** Plot of 2^nd^ pulse-evoked peak amplitudes (mean ± SEM) against hyperpolarizing gap duration. **e:** Plot of channel availability against pre-pulse potential. **f:** Plot of 2^nd^ pulse-evoked amplitudes against 1^st^ pulse duration.

Covalent binding-induced changes in channel gating kinetics (**RFI** and **SDO**) and gating equilibrium (**SSI**) are shown in Figures 5b,c,d. Blue, teal, and green colors indicate resting-state-, V-half-, and inactivated-state-illumination, respectively. Insets show normalized data, where amplitudes were normalized to the maximal amplitude after azido-riluzole perfusion and irradiation. This gives a clearer picture of how the gating has been modulated. Similarly to our earlier data^9^ obtained using continuous illumination, the population of ion channels that were still conducting showed modulated gating: delayed recovery from inactivation (Fig. 5b), shifted steady-state availability (Fig. 5c), and accelerated onset of inhibition (Fig. 5d). Note that the **SDO** curves were also affected by covalently bound ligand; in the case of covalently bound azido-riluzole after the inactivated-state-illumination protocol, the onset proceeded with a time constant of 1.21 ± 0.12 ms.

Since there was no possibility of binding or unbinding, this must correspond with a conformational rearrangement of the ligand-channel complex, which was somewhat slower than in the case of 100 μM riluzole (τ = 0.35 ms)^9^.

Interestingly, 90 ms UV pulses delivered at every 400 ms during inactivated conformation were as effective as continuous illumination had been^9^, in spite of the fact that the total illumination time was only 23.75% of it. It seems that repeatedly allowing azido-riluzole molecules to diffuse to their most favorable location (probably the binding site) without activating them, and then delivering the UV pulse only when they are at the right location, actually works as well as continuous illumination.

In contrast, when UV pulses were delivered during resting state, no significant decrease of current amplitudes was observed, and changes in recovery kinetics (**RFI** protocol) and equilibrium of inactivation (**SSI** protocol) were also non-significant (p = 0.2 and p = 0.25 for changes in the predominant time constant of recovery and V_1/2_, respectively, n = 7). This may either suggest that the central cavity or the binding site itself is not accessible at resting conformation, or that the binding site has very low affinity at resting state. UV pulses delivered at the approximate half-inactivation voltage were between the other two illumination protocols in effectiveness.

#### Moderate illumination intensity reveals the relative contribution of modulation and channel block

In the experiments described thus far, we intended to investigate the case when the local anesthetic binding site is fully saturated, i.e., the whole channel population binds at least one azido-riluzole molecule. (Close to full saturation is shown by the fact that in the **RFI** protocol at gap durations <1 ms currents were almost fully inhibited – see Fig. 5b). However, if we want to study a channel population with full occupancy of binding sites, nonspecific binding to other sites would unavoidably occur. Binding to multiple binding sites would then render the channels nonconducting. In this series of experiments we intended to investigate the contribution of non-blocking modulation in a situation that resembles therapeutic concentrations: when specific binding is predominant, and nonspecific binding is minimal. In order to do this, we decreased the intensity of the UV light by moving the light source somewhat further away from the measured cell (5-7 mm above the cell, instead of 3-4 mm), and re-designed the voltage protocol that was being executed during azido-riluzole perfusion and pulsed UV illumination (Fig. 6a). The 90 ms UV light pulses were delivered at depolarized membrane potential (like in the case of “inactivated-state-illumination"), but three extra depolarizations were included afterwards, in order to monitor the onset of drug-induced modulation of gating. The 1^st^ depolarization served as control, and at the same time it was used to deliver the UV pulse. The 2^nd^ and 3^rd^ depolarizations were meant to assess recovery kinetics, while the 4^th^ one to assess the inactivation/recovery equilibrium. During experiments, we monitored the current evoked by the 4^th^ depolarization. When it was inhibited by ~80%, pulsed illumination was stopped and unbound azido-riluzole was washed out. As it is shown in Fig. 4c, at this point the current evoked by the 1^st^ depolarization was only inhibited to 92.2 ± 3.4% of the original amplitude. Currents evoked by 2^nd^, 3^rd^, and 4^th^ pulses were inhibited to 79.6 ± 2.9%, 51.9 ± 3.9%, and 17.6 ± 1.2% of their respective control amplitude. At this point, cells were again tested by **RFI**, **SSI** and **SDO** protocols (Fig. 4d,e,f; red lines). In spite of the minimal decrease in 1^st^ pulse-evoked amplitudes, these protocols showed that specific binding sites were already occupied by covalently bound ligands: the patterns of inhibition observed in **RFI** and **SDO** protocols were very similar to the patterns observed after using the “inactivated-state-illumination” protocol. Interestingly, the shift of the **SSI** curve was much less than in the case of the “inactivated-state-illumination” protocol, indicating that binding of further molecules to nonspecific sites can add to the shift of equilibrium steady-state inactivation. What fraction of the channel population had its local anesthetic binding site occupied after the photolabeling? We can tell this from the maximal inhibition caused by the covalently bound ligand. The inhibition was maximal at the shortest gap durations, however at very small currents accurate measurement was hindered by the poor signal to noise ratio. At 0.66 ms hyperpolarizing gap length, where the signal to noise ratio was already acceptable (Fig. 6c), the current was inhibited to 14.1 ± 2.4 % of the control, while at 66 ms gap length it was only inhibited to 85.5 ± 5.7 % of the control. Note that both inhibition values were caused by the exact same fraction of occupied binding sites. If we suppose that the 85.5 % at 66 ms represented unblocked channels, then >83.7 % of this unblocked fraction had to be inhibited by modulation to reach the inhibition to <14.1 % at short gap lengths (0.855 * (1 - 0.837) = 14.1). We can conclude that non-blocking modulation alone can cause at least 83.7 % inhibition in the case of covalently bound azido-riluzole.

**Fig. 6.**
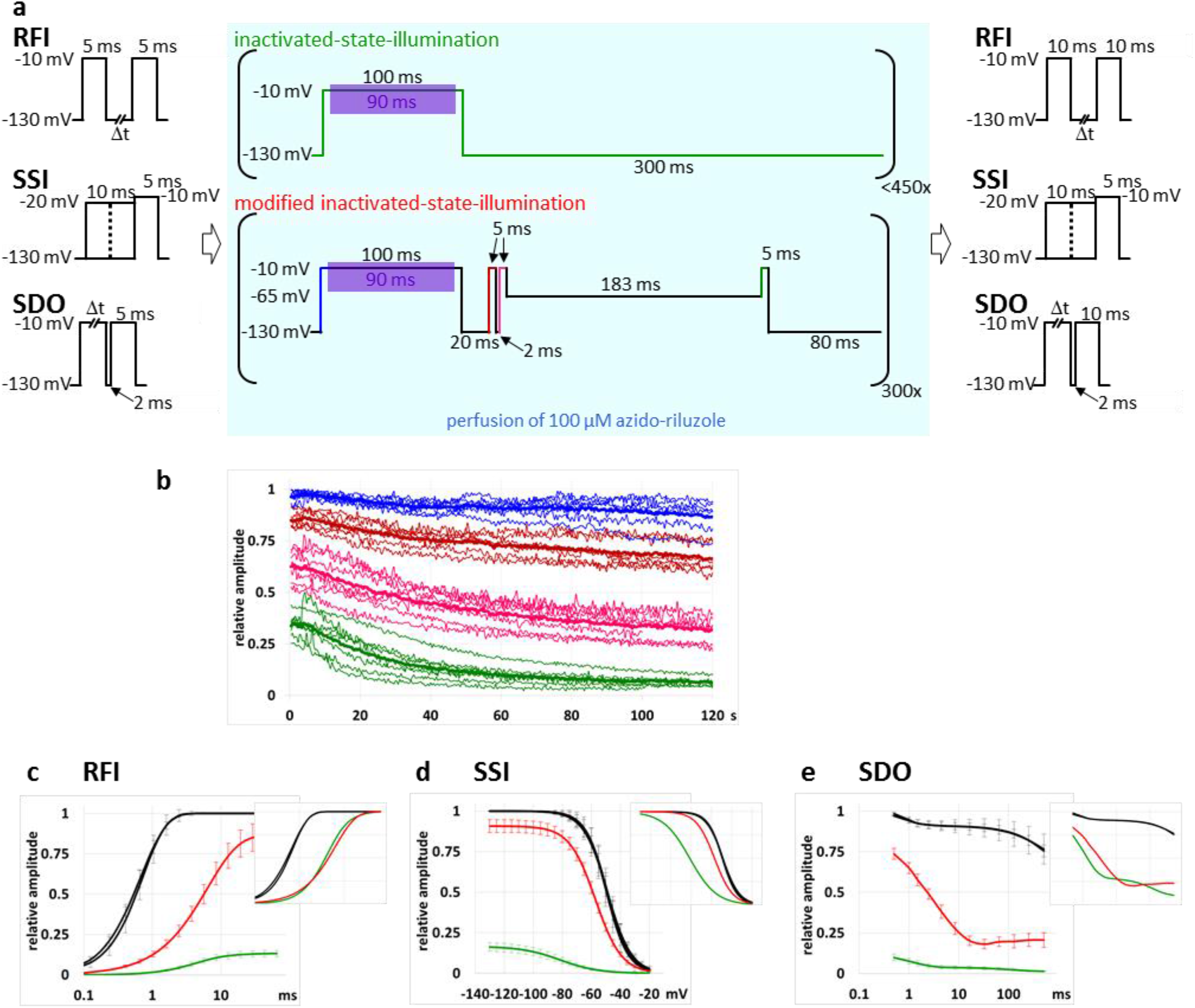
Maximizing modulation and minimizing channel block in inactivated-state-illumination experiments with decreased intensity. **a** Experimental protocol. **RFI**, **SSI**, and **SDO** protocols were run before and after azido-riluzole perfusion and pulsed UV illumination. The inactivated-state-illumination protocol is the same as in Fig. 5. The modified protocol contains three additional depolarizations to monitor changes in gating kinetics and equilibrium. The color of depolarizations in the voltage protocol is the same as the color of plots in panel **b**. **b** The plot of 1^st^ (blue), 2^nd^ (dark red), 3^rd^ (magenta) and 4^th^ (green) pulse-evoked current amplitudes in n = 7 cells, normalized to the 1^st^ pulse-evoked amplitude at the start of the experiment. Thick lines indicate averages. **c, d, e** Assessment of gating kinetics and equilibrium before and after azido-riluzole perfusion and UV irradiation. Black lines show control data before azido-riluzole perfusion and UV illumination, colored lines show data measured after stopping UV pulses and washing out azido-riluzole. Green lines are shown for comparison and are identical to the data in Fig. 5; red-lines show data after the modified inactivated-state-illumination protocol. Amplitudes were normalized to the maximal amplitude in control. Insets: Amplitudes normalized each to its own maxima. **d:** Plot of 2^nd^ pulse-evoked peak amplitudes (mean ± SEM) against hyperpolarizing gap duration. **e:** Plot of channel availability against pre-pulse potential. **f:** Plot of 2^nd^ pulse-evoked amplitudes against 1^st^ pulse duration.

#### Specificity of binding

Next, we investigated how specific the binding was, by testing the F1579A binding site mutant channels. We expected, that if the binding was indeed specific, then we would see neither inhibition of the amplitude, nor modulation of gating. Interestingly, mutation of the binding site did not prevent inhibition of the amplitude (provided that channels assumed inactivated conformation), but modulation of gating was radically decreased (Fig. 7a,b,c). This suggests that when the high affinity binding site is disrupted, inhibition is still possible, probably by binding to secondary binding sites, but these binding sites are less potent in conferring gating modulation. The involvement of this phenylalanine in the mechanism of modulation has been shown before^13,14,16–19^, therefore the lack of significant modulation is not unexpected. However, the fact that mutant channels could still be effectively inhibited in inactivated state, was indeed unexpected, and it suggests the existence of one or more additional binding sites. Association to these binding sites, which were only available in the inactivated conformation, caused predominantly channel block, not modulation.

**Fig. 7.**
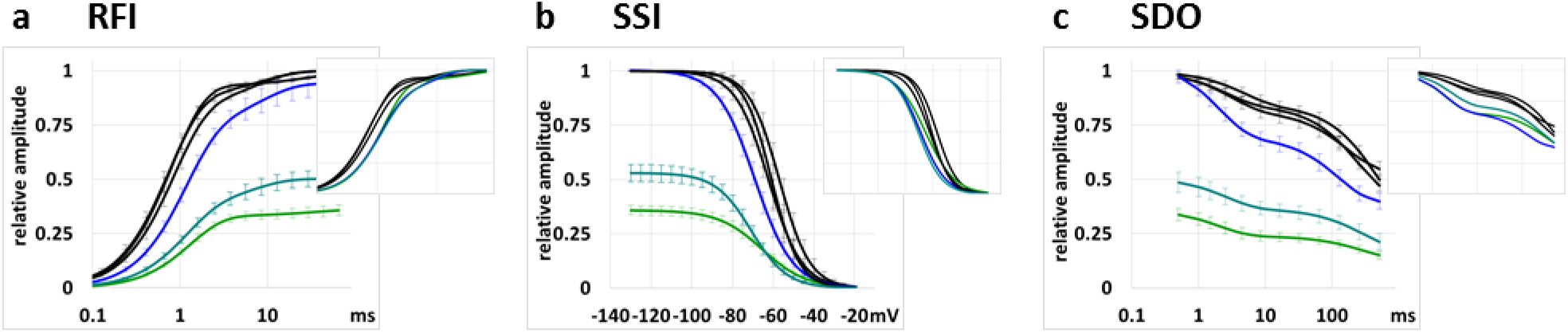
Effect of conformation-selective photolabeling by azido-riluzole on gating kinetics of F1579A mutant channels. **a, b, c** Assessment of gating kinetics and equilibrium before and after azido-riluzole perfusion and UV irradiation. Black lines show control data before azido-riluzole perfusion and UV illumination, colored lines show data measured after stopping UV pulses and washing out azido-riluzole. Blue lines: after resting-state-illumination protocol; teal lines: after V-half-illumination protocol; green lines: after inactivated-state-illumination protocol. Amplitudes were normalized to the maximal amplitude in control. Insets: Amplitudes normalized each to its own maxima. **a:** Plot of 2^nd^ pulse-evoked peak amplitudes (mean ± SEM) against hyperpolarizing gap duration. **b:** Plot of channel availability against pre-pulse potential. **c:** Plot of 2^nd^ pulse-evoked amplitudes against 1^st^ pulse duration.

#### Can channels recover from inactivation without unbinding?

From the experiments discussed thus far we can conclude that the fast recovery process can be observed even when there was no dissociation, therefore, it is predominantly due to modulated gating. This fast recovery process was practically complete within ~10 ms in the case of either riluzole or covalently bound azido-riluzole, and within ~3 ms in the case of unactivated azido-riluzole (see Fig. 4c in ref^9^). In our next experiment we aimed to test if unbinding contributes to either of the recovery processes. We reasoned that if the fast process indeed did not involve unbinding as we supposed, then we could catch inhibitor molecules undissociated and bind them covalently, even after channels recovered from inactivated state. To test this possibility, we changed the timing of UV pulses, delivering pulses 5 ms after stepping back to the holding potential (Fig. 8a). We found that inhibition, and – more significantly – modulation were almost as effective as in the inactivated-state-illumination protocol (Fig. 8b,c,d). This indicates that azido-riluzole molecules must have stayed effectively bound (being close enough, and in similar orientation) for at least a few tens of milliseconds after the channel had regained the resting conformation, because they could form a covalent bond that resulted in the exact same type of modulation. In contrast, by the end of a 205 ms period spent in resting conformation, azido-riluzole molecules apparently had become unbound and therefore effective covalent binding to the binding site could not occur. The reason for this may be that molecules had already left the central cavity, or that the binding sites had become unavailable (e.g. buried by a twist of the S6 helix). Note that these data reflect the dynamics of azido-riluzole binding/unbinding, which is different from that of riluzole. We have no means to directly test how long riluzole stays at the binding site after resting conformation has been regained. However, considering that azido-riluzole has an approximately tenfold lower affinity to sodium channels (without UV irradiation 100 μM azido-riluzole was roughly equieffective with 10 μM riluzole^9^), it is safe to assume that unbinding of riluzole cannot be faster than that of azido-riluzole, and thus cannot contribute to the fast recovery process. We suppose that it is most likely to form the early component of the slow recovery process (with time constants ranging from 50 to 190 ms, see Fig. 2g and Supplementary Fig. 1), because that would be consistent with the fast association rate (τ < 4.81 ms^20^), and the calculated affinity values to inactivated state (0.29 μM^21^, 0.2 μM^22^, or 0.3 μM^23^). We suppose that the late phase (~750 ms) most probably reflects depletion of intracellular stores, and partitioning into extracellular aqueous phase, which may be why this component was also observed in experiments with binding site mutant channels.

**Fig. 8.**
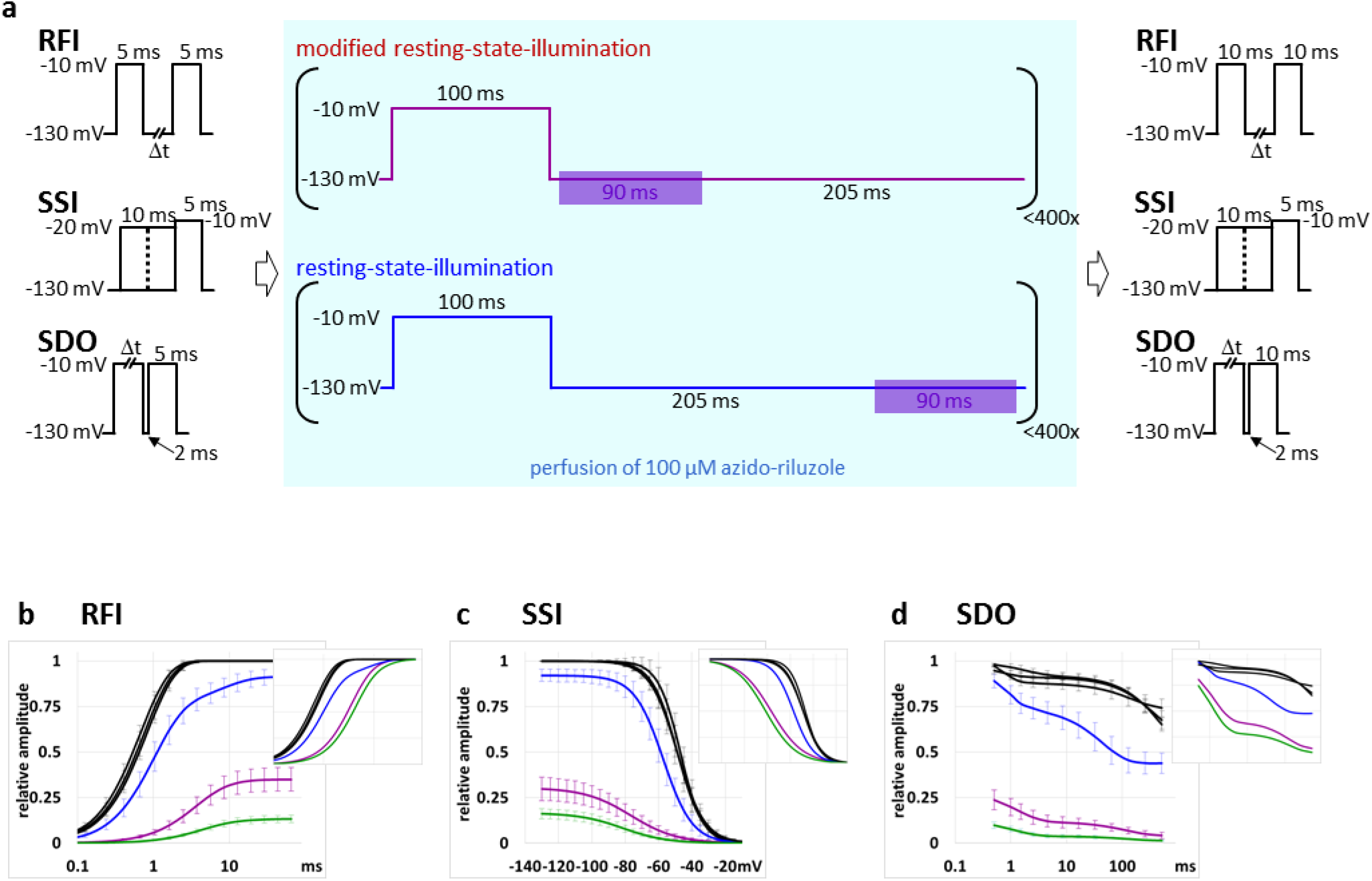
Timing of UV pulses during hyperpolarization reveals unbinding dynamics of azido-riluzole. **a** Experimental protocol, as described before. The resting-state-illumination protocol is shown for comparison. **b, c, d** Assessment of gating kinetics and equilibrium before and after azido-riluzole perfusion and UV irradiation. Black lines show control data, purple lines show measurements after the modified resting-state-inhibition. Measurements after resting-state-illumination (blue lines) and inactivated-state-illumination (green) are shown for comparison; they are identical with the data shown in Fig.5. **b:** Plot of 2^nd^ pulse-evoked peak amplitudes (mean ± SEM) against hyperpolarizing gap duration. **c:** Plot of channel availability against pre-pulse potential. **d:** Plot of 2^nd^ pulse-evoked amplitudes against 1^st^ pulse duration.

### Possible location of bound riluzole

In order to gain insights into the possible binding sites of riluzole, we performed *in silico* docking of this small molecule to the wild type and mutant Na_V_1.4 structures. The human Na_V_1.4 (PDBID: 6AGF) was used, since *in silico* docking is more reliable to experimental structures than to homology models. Importantly, riluzole binding was not observed in the pore region or close to the pore in the central cavity in either the wild type or the F1586A (F1579A in rNa_V_1.4) structures (Fig. 9). This inhibitor bound among transmembrane helices in three fenestrations of the wild type structure. In all cases, interactions involved the aromatic side chain of phenylalanine residues (F436, F1284, and F1586; which correspond to F430, F1277, and F1579 in rNa_V_1.4) are indicated. In the absence of the phenylalanine side chain in the F1586A mutant, only two binding sites were observed at F436 and F1284.

**Fig. 9.**
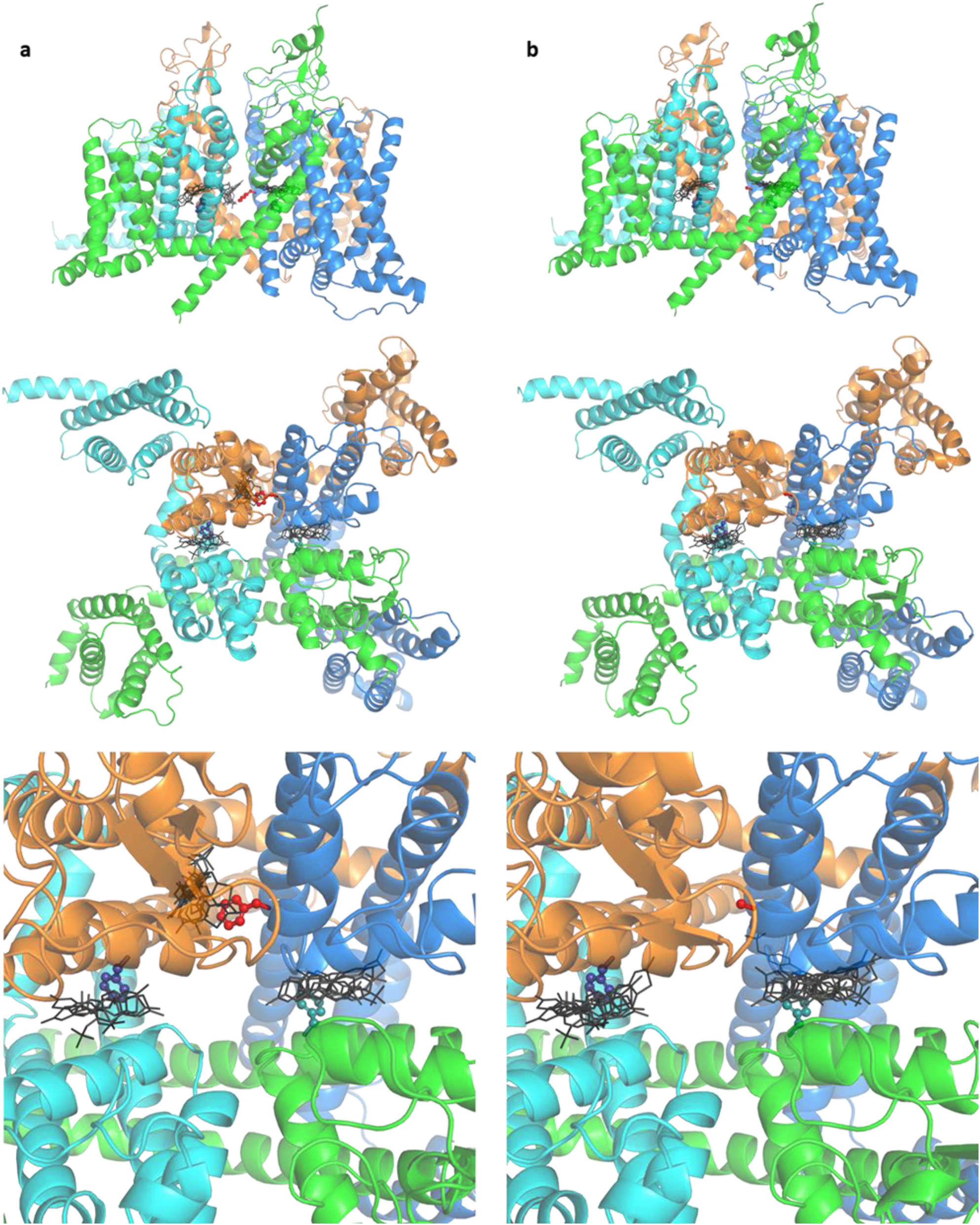
Riluzole does not block the pore. Riluzole was docked into the structural model of human Na_V_1.4 (PDB ID: 6AGF) (**a**) and its F1586A mutant (**b**). Gray sticks: best poses of riluzole; sticks and balls: F436 (teal), F1284 (blue), and F1586 (red); green, cyan, orange and marine: domains I, II, III and IV.

**Fig. 10.**
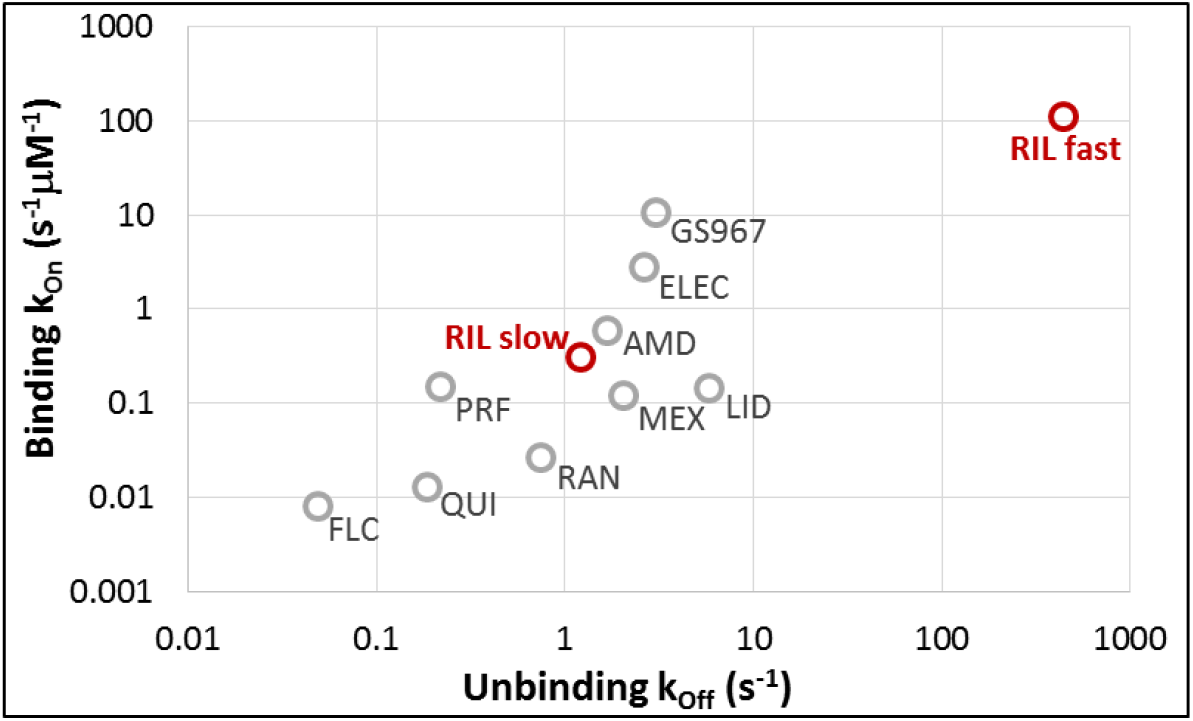
Comparison of calculated binding and unbinding rates of riluzole to 9 sodium channel inhibitor compounds. The figure was re-plotted from El-Bizri et al., 2018^64^., and supplemented with our data with riluzole. (AMD–amiodarone; ELEC – eleclazine; FLC – flecainide; LID – lidocaine; MEX– mexiletine; PRF – propafenone; QUI – quinidine; RIL – riluzole) Binding and unbinding rates were calculated for time constants of both the fast and the slow recovery processes.

## Discussion

Our data suggest that riluzole is the first member of an entirely new class of sodium channel inhibitor compounds. Its primary binding site is the well-characterized “local anesthetic receptor", with F1579 as the key residue. However, both the location of the bound ligand, and the effect exerted by it are radically different. As for the location, *in silico* docking revealed that unlike other sodium channel inhibitor molecules, riluzole was preferentially located within the fenestrations, and tended to avoid the central part of the pore or the vicinity of the selectivity filter, where it could interfere with conduction. Localization within the fenestration has been previously observed in molecular dynamics simulations investigating the general anesthetic isoflurane^24^ and propofol^25^, as well as the local anesthetic benzocaine^26–28^ and lidocaine (neutral form)^29^, but it has never been found to be the dominant position. The unusual location predicted by in silico docking is supported by the way riluzole acted in experiments. It stabilized inactivated conformation, without interference with conduction. Inactivated state stabilization affected both the equilibrium and the recovery rate as evidenced by the shift of the **SSI** and the **RFI** curves, respectively (Fig. 1; see also Fig. 1 in ref.^9^, and Fig.2 in ref^10^). In addition, in the **SDO** protocols, we observed that onset of inhibition by riluzole was not instantaneous (it occurred with τ = 4.89 ± 0.53 ms at 10 μM, and τ = 0.35 ± 0.056 ms at 100 μM)^9^. Interestingly, this non-instantaneous onset was observed even with covalently bound azido-riluzole (τ = 0.68 ± 0.09 ms), suggesting that it is based on a conformational rearrangement of the channel-ligand complex. In this study we found similar non-instantaneous onset (τ = 1.21 ± 0.12 ms in the “inactivated-state-illumination” protocol).

The non-instantaneous onset (as seen in the **SDO** protocol) and the delayed recovery (see the **RFI** protocol) together ensure that effective inhibition is confined within a strict temporal window. Single pulse-evoked currents, or synaptic activity-evoked action potentials are “missed” by riluzole, because inhibition becomes effective only after the channel has reached inactivated conformation, and the conformation of the ligand-channel complex has been stabilized. After this, inhibition is effective, but only for a limited time – even in the case of the non-therapeutic concentration of 100 μM, at the non-physiological membrane potential of −130 mV it was only effective for approximately 10-20 ms. At therapeutic concentrations the effect of riluzole is no more than a fine modulation: the refractory time after an action potential is prolonged by a few milliseconds, and a small fraction of the channel population is kept in inactivated conformation. The exact extent of prolongation and of the fraction kept in inactivated conformation is dependent on the membrane potential during the interspike interval. This temporal window of effectiveness is the basis of both persistent (or “late") sodium current selectivity, and the low pass filtering effect. The contribution of the persistent component of sodium currents (INaP) is significant during gradual depolarizations preceding action potentials, especially during burst firing^30–33^. In these cases there is time enough for the onset of inhibition, and the inhibition remains effective until the membrane becomes hyperpolarized for a sufficient time.

Upregulation of the persistent sodium current has been shown to be involved in a number of pathologies, including epilepsies^34,35^, cardiac pathologies^3,36–41^, neurodegenerative disorders^42–46^, pain disorders^47^, even cancer metastasis^48^ and type II diabetes^49^. Selective inhibitors of I_NaP_ have been found to be effective in cardiac diseases^38,41,50–52^, epilepsies^53–55^, pain syndromes^56–58^, nerve injuries and neurodegenerative diseases. Riluzole has been found to be especially effective in preventing damage in spinal cord and peripheral nerve injury^59–62^. Riluzole represents an entirely new class among I_NaP_ selective inhibitors, because the non-blocking modulation mechanism endows it with an “ultrafast” offset kinetics, which, however is not based on actual dissociation. Similar mechanism has been assumed for inhibition by riluzole in a recent computational study^63^. To illustrate how this compares to offset kinetics of other well known sodium channel inhibitors, we re-plotted the data from a comparative study of nine inhibitor compounds^64^ (see Table 2 and Fig. 4D of the quoted paper), which included six antiarrhythmic drugs, and the three best known I_NaP_ selective inhibitors. We supplemented this figure with our data from experiments with riluzole (Fig. 8). In the quoted paper binding rate constants were plotted against unbinding rate constants. The latter was calculated from the time constants of the offset (*k_off_ = 1 / τ_off_*), then the former was calculated from *k_off_* and the *IC_50_* value (*k_on_ = k_off_ / IC_50_*). We calculated *k_on_* and *k_off_* values the same way for riluzole using time constants of both the fast and slow recovery processes (τ_fast_ = 2.25 ms, and τ_slow_ = 823 ms), and the IC_50_ value 3.98 μM (from inhibition of 3^rd^ pulse-evoked currents in the **3PT** protocol). It is clear that as regards the slow recovery process (which, at least in part, reflects true dissociation) riluzole did not differ from the other compounds. However, riluzole has an additional “power": It can “pretend” to have been dissociated, which practically increases its speed 366-fold. This allows it to be effective in frequency ranges that are unavailable for conventional drugs. This in fact has already been observed in several experimental studies, where riluzole has been found to be especially potent at high frequencies (see *e.g.* Table 2 in Desaphy *et al.*^65^, Fig. 5 in Urbani and Belluzzi^32^).

Note that *IC_50_* and *k_off_* data predict that *k_on_* should be extremely high (112 s^−1^μM^−1^), which, using the equation: *τ_on_* = 1/(*cc*\**k_on_* + *k_off_*), would give *τ_on_* = 86 μs at *cc* = 100μM concentration – which is clearly contrary to experimental data.

To further illustrate how riluzole differs from other sodium channel inhibitors we used the data from Table 2 in Desaphy *et al.*^65^, which gives *IC_50_* values for six drugs measured at sodium currents evoked at 0.1 Hz or 10 Hz from −120 mV holding potential, as well as at 50 Hz from −90 mV holding potential. We calculated the ratios of *IC_50_* values and plotted the 0.1 Hz / 10 Hz ratios against the 10 Hz / 50 Hz ratios (Fig. 11). It is clear from this representation that riluzole is in a different class of inhibitors. Most drugs showed higher *in vitro* potency when the frequency of depolarizations was increased from 0.1 Hz to 10 Hz, but did not change much between 10 and 50 Hz. Riluzole, on the other hand, only started to “realize its potential” above 10 Hz, the apparent affinity increased 48-fold, from *IC_50_* = 43 μM (10 Hz) to *IC_50_* = 0.9 μM (50 Hz).

**Fig. 11.**
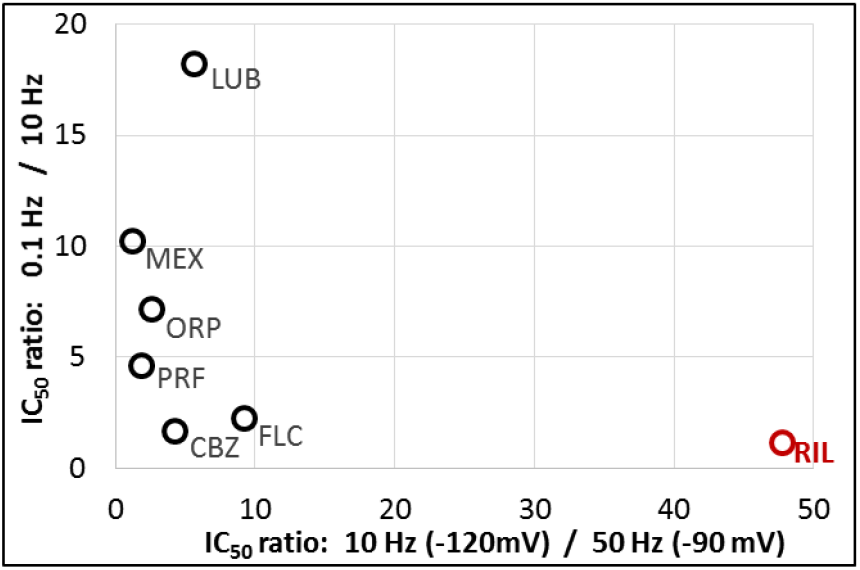
Differences in the frequency-dependence of inhibition between riluzole and 6 other sodium channel inhibitor compounds. We used data from Desaphy et al., 2014^65^., calculated ratios of apparent affinities (IC50 values), and plotted increase of apparent affinity between 0.1 Hz and 10 Hz against increase of apparent affinitiy between 10 Hz and 50 Hz. (CBZ – carbamazepine; FLC – flecainide; LUB – lubeluzole; MEX – mexiletine; ORP – orphenadrine; PRF – propafenone; RIL – riluzole)

In summary, our data indicate that riluzole is able to bind with fast onset kinetics, and high affinity to the conventional local anesthetic binding site of sodium channels, but only in open or inactivated conformations. The binding itself does not prevent conduction, only slightly alters the gating of channels, making them more likely to stay in inactivated conformation. This produces a preferential inhibition of i) I_NaP_, ii) cells with depolarized membrane potential, and iii) cells firing at high frequencies. The combination of increased persistent component, compromised ability to keep resting membrane potential, and abnormally high firing frequency almost always signifies pathology, which is the reason why riluzole can selectively inhibit this type of pathological activity. The basis of this selectivity is the non-blocking modulation mechanism. We anticipate that compounds of this class (fast acting non-blocking modulators) may be favorable for the treatment of a number of pathologies, such as neuropathic pain syndromes, certain neuromuscular diseases, epilepsies, and cardiac fibrillations, which are characterized by an enlarged persistent sodium current and high frequency firing.

## Methods

### Cell culture and expression of recombinant sodium channels

Cloning of the wild type and F1579A mutant rNa_V_1.4 channel constructs was done as described previously^9^. Recombinant sodium channel expressing cell lines were generated by the transfection of wild type and F1579A mutant Na_V_1.4 BAC DNA constructs into HEK 293 cells (ATCC CRL-1573) by Fugene HD (Promega, Fitchburg, WI) transfection reagent according to the manufacturer’s recommendations. Cell clones with stable vector DNA integration were selected by the addition of Geneticin (Life Technologies, Carlsbad, CA) antibiotic to the culture media (400 mg/ml) for 14 days. HEK293 cells were maintained in Dulbecco’s Modified Eagle Medium, high glucose supplemented with 10% v/v fetal bovine serum, 100 U/ml of penicillin/streptomycin and 0.4 mg/mL Geneticin (Life Technologies, Carlsbad, CA). For experiments cells were plated onto 35 mm Petri dishes or T75 flasks, and cultured for 24–36 hours. For manual patch clamp experiments Petri dishes were transferred to the recording chamber, where cells were kept under continuous flow of extracellular solution. For Port-a-Patch experiments cells were dissociated from the dish with trypsin-EDTA, centrifuged and suspended into the extracellular solution.

### Materials

All chemicals were obtained from Sigma-Aldrich. Azido-riluzole was synthesized by SONEAS Research Ltd., Budapest, Hungary.

### Electrophysiology

The composition of solutions (in mM) was: Intracellular solution: 50 CsCl, 10 NaCl, 60 CsF, 20 EGTA, 10 HEPES; pH 7.2 (adjusted with 1 M CsOH). Extracellular solution: 140 NaCl, 4 KCl, 1 MgCl_2_, 2 CaCl_2_, 5 D-Glucose and 10 HEPES; pH 7.4 (adjusted with 1 M NaOH). Osmolality of intra- and extracellular solutions was set to ~290 and ~300 mOsm, respectively. Data were sampled at 20 kHz, and filtered at 10 kHz. Experiments were carried out at room temperature.

In manual patch clamp experiments whole-cell currents were recorded using an Axopatch 200B amplifier and the pClamp software (Molecular Devices, San Jose, CA). Pipette resistances ranged between 1.5 and 4.0 MΩ. Solution exchange was performed by the “liquid filament switch” method^66,67^, using a Burleigh LSS-3200 ultrafast solution switching system, as described in detail previously^68^. Flow rate was set using a DAD-12 solution exchange system (ALA Scientific Instruments Inc., Farmingdale, NY), to 0.2-0.3 ml/min, which corresponded to 5-20 cm/s velocity. Ten to 90% solution exchange rates were between 1 and 3 ms. In experiments with the Port-a-Patch system (Nanion, Munich, Germany) currents were recorded using an EPC10 plus amplifier, and the PatchMaster software (HEKA Electronic, Lambrecht, Germany). During cell catching, sealing, and whole-cell formation, the PatchControl software (Nanion) commanded the amplifier and the pressure control unit. The resistance of borosilicate chips was 2.0–3.5 MΩ.

### UV photoactivation

The recording chamber of the Port-a-Patch was customized to accommodate a 400 μm diameter quartz optic fiber, which was placed 3–4 mm (or 5-7 mm in a subset of experiments) above the recorded cell. The original perfusion manifold was replaced by a custom manifold positioned to the side of the recording chamber opposite to the waste removal. Solution exchange was complete within 1–2 s. UV light pulses were delivered by a 310 nm fiber coupled Mightex FCS-0310-000 LED (Mightex, Pleasanton, CA), with 40 μW intensity. Pulses were triggered by the PatchMaster software.

### Data analysis

Curve fitting was done in Microsoft Excel, using the Solver Add-in. Steady-state inactivation (**SSI**) curves were fitted using the Boltzmann function: *I* = *I_max_*/{1 + exp[*V_p_* – *V_1/2_*/ -*k*]}, where V_p_ is the pre-pulse potential, V_1/2_ is the voltage where the curve reaches its midpoint, and k is the slope factor.

Recovery from inactivation (**RFI**) experiments were fitted by a mono-, or bi-exponential function: *I* = *A_1_*\*[1-exp(*-t_ip_*/*τ_1_*)]^*n*^ + *A_2_*\*[1-exp(*-t_ip_*/*τ_2_*)], where *τ_1_* and *τ_2_* are the fast and slow time constants, *A_1_* and *A_2_* are their contributions to the amplitude (*A_2_* = 0, for monoexponential fitting), and *t_ip_* is the duration of the interpulse interval. We observed that recovery in the presence of riluzole was steeper than a simple exponential function, therefore we had to introduce the exponent *n*. When its value was unconstrained, best fits produced *n* = 2.35 ± 0.65, range: 1.22 to 3.15. However, the value of the exponent affected the value of the time constant, as we have discussed before^9^. For this reason we chose *n* = 1.5, the smallest exponent that could acceptably fit our data, and used it for fitting throughout the experiments.

State-dependent onset (**SDO**) data were fitted with either single or double exponential functions: *I* = *A1*\*exp(*-t_p_*/*τ_1_*) + *A_2_*\*exp(*-t_p_*/*τ_2_*), where *A_1_* and *A_2_* are the relative amplitudes of the two components, and *t_p_* is the duration of the pulse.

Averaging of **SSI**, **RFI** and **SDO** curves was not done by calculating the mean of measured points, because this would introduce an error in the slope of curves. Instead, experimental curves were individually fit, and the parameters were averaged (arithmetic mean for amplitudes, membrane potential values, and slope factors; geometric mean for time constants), then a curve was re-plotted using the averaged parameters.

Apparent affinity (*IC_50_*) values were calculated either from the extent of inhibition or from onset and offset time constants. Apparent affinity, as we have discussed before^10^, is a useful way to represent the potency of sodium channel inhibitors under specific circumstances. The potency can change radically even on the sub-millisecond time scale, as seen for example in the **3PT** protocol, depending on the temporal pattern of membrane potential, due to the interrelated dynamics of binding and gating. Apparent affinity values can be calculated from a single inhibition value, using the simplified Hill equation: when one-to-one binding is assumed, the Hill equation is reduced to *Inh* = *cc*/(*cc* + *IC_50_*), where *Inh* is the inhibited fraction and *cc* is the drug concentration. The calculation is most accurate at ~50 % inhibition, but becomes increasingly inaccurate as inhibition approaches either 0 or 100 %. The *IC_50_* value can also be calculated from onset and offset time constants (obtained by single exponential fitting of amplitudes in the **3PT** protocol), if we suppose a single-step binding reaction: *IC_50_* = *cc*\* *τ_on_*/(*τ_off_* -*τ_on_*). In the case of riluzole, the inhibited fraction and the bound fraction are not equivalent, therefore we expect that the IC_50_ values calculated by the two ways will differ (for the results see Supplemental Fig. S1).

Peak amplitudes from the 3PT protocol showed a slow decrease during experiments (see Fig. 2, 4 and Supplemental Fig. S1), due to slow inactivation (more exactly to a fast-entry-slow-exit form of inactivation^69,70^). In order to appropriately calculate the extent of inhibition by riluzole, we corrected for the slow inactivated fraction: A sum of an exponential and a linear component was fit to the first 6 and last 3 points of 1^st^ pulse-evoked current amplitudes in each cell (Fig. 2e, 4e, and Supplemental Fig. S1b and h). This fitted function represented the non-slow-inactivated fraction *f_nsi_(t)* of the channel population. Measured amplitude plots (*A_M_(t)*), then were transformed into corrected amplitude plots (*A_C_(t)*) by expressing it as a fraction of non-slow-inactivated channels: *A_C_(t) = A_M_(t) / f_nsi_(t)*.

### Structural models and in silico docking

The human Na_V_1.4 (PDBID: 6AGF) was used, since *in silico* docking is more reliable to experimental structures than to homology models. The F1586A mutant structure was prepared manually by removing atoms of Phe and renaming the residue to Ala. The protein and riluzole (ZINC database ID: ZINC26671469) structures were prepared using prepare_ligand4.py and prepare_receptor4.py scripts from MGLTools with default options except hydrogens were added in the case of the protein (https://ccsb.scripps.edu/mgltools). Riluzole was docked to the wild type and F1586A mutant structures using Autodock Vina^71^. The search space was setup in PyMOL (The PyMOL Molecular Graphics System, Version 2.4, Schrödinger, LLC.) via the Autodock/Vina plugin^72^ (Supplemental Fig. S2). The exhaustiveness option of Vina was set 128 instead of the default 8 for increasing the probability of finding the minimum of the scoring function. The num_modes and energy_range options were set to 100 and 20, respectively, to output a higher number of docked poses.

---

**Fig. S1.**
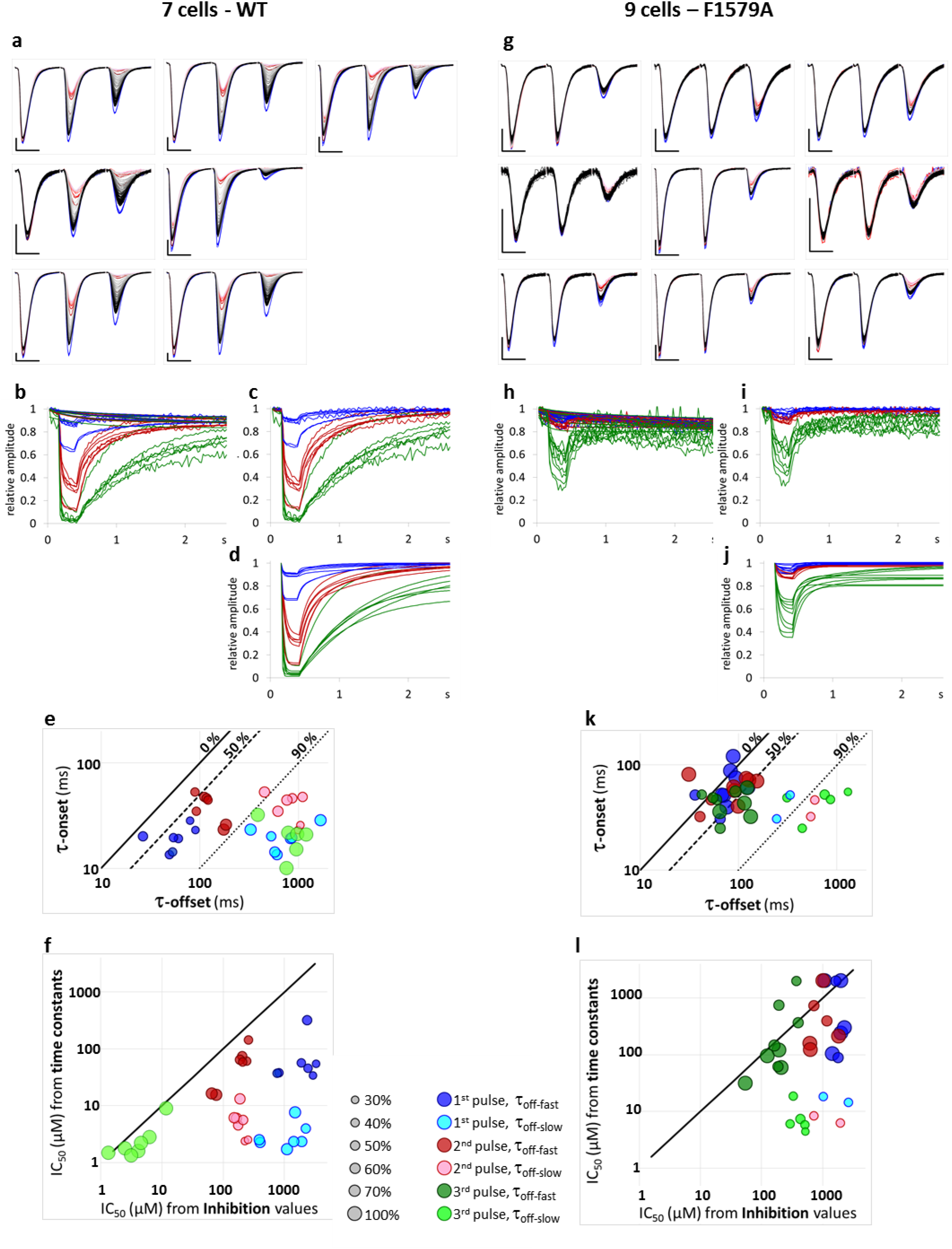
Calculation and evaluation of onset and offset time constants. **a:** The 1^st^, 2^nd^, and 3^rd^ pulse-evoked currents recorded in 7 cells throughout the **3PT** experiment, before (blue traces), during (dark to light red traces), and after (light gray to black traces). Scale bar: 1 nA, 1 ms. **b** Peak amplitude plots for the currents shown above. Each of the three pulse-evoked currents are normalized to its own control. The effect of slow inactivation was estimated by fitting the sum of an exponential and a linear component to the first 6 and last 3 points of 1^st^ pulse-evoked current amplitudes in each cell (shown by thin lines). **c** The same traces after correction for slow inactivation. **d** Exponentials fitted to the curves shown in panel c. **e** Onset time constants plotted against offset time constants. Wherever the offset was fit by a biexponential function, fast and slow components are shown by dark and light colors, respectively, the size of circles indicating their respective contribution. (Third pulse-evoked amplitudes were adequately fitted with monoexponential function in each cell.) If one supposed a single-step binding reaction, the ratio of onset and offset time constants would correspond with the extent of inhibition. For example 90 % inhibition would correspond with a tenfold difference between onset and offset rates (marked by a dotted line), a 50 % inhibition would correspond with a twofold difference (dashed line), and approaching minimal inhibition the onset rate would approach the offset rate (solid line). It is clear that time constants did not correctly reflect the extent of observed inhibition, for example 1^st^ pulse-evoked currents were inhibited by 15.2 ± 3.6 %, while fast and slow time constants indicated >50 % and >90 % inhibition, respectively. **f** Apparent affinity (IC50) values calculated from the inhibited fraction were plotted against IC50 values calculated from time constants (see Methods). Time constants clearly revealed higher apparent affinity, indicating that a large part of binding site occupancy did not translate into effective inhibition. Size and color of circles indicate contribution of exponential components as in panel **e**. **g** to **l** The same data as in panels **a** to **f**, for n = 9 cells expressing F1579A mutant channels. Offset was monoexponential in 7 out of 9 cells in the case of 1^st^ and 2^nd^ pulse-evoked current amplitudes, and in 4 out of 9 cells in the case of 3^rd^ pulse-evoked current amplitudes. In the rest of the cells a slow component was also detectable. Note that the difference between inhibition-derived IC50 values and time constant-derived IC50 values was much less than in WT channels, suggesting that a much larger part of binding site occupancy translated into inhibition, *i.e.*, non-blocking inhibition had a minor contribution.

**Fig. S2.**
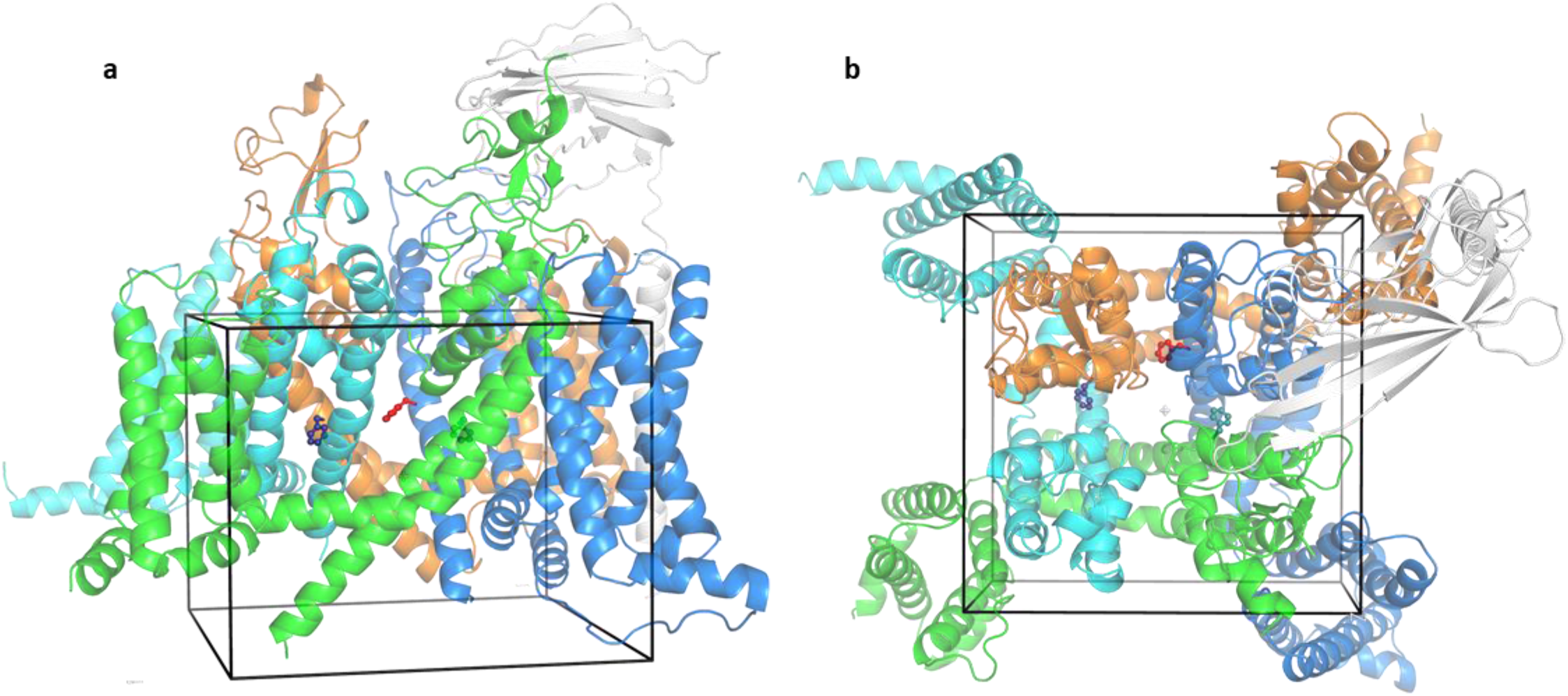
The search space for riluzole docking. The box (black) was defined in PyMOL using Autdock/Vina plugin and included the central cavity, the cytosolic region below this cavity and a region of the selectivity filter. Side (**a**) and top (**b**) views of the protein and box are shown. Sticks and balls: F436 (teal), F1284 (blue), and F1586 (red); green, cyan, orange and marine: domains I to IV of Na_V_1.4 subunit alpha; white cartoon: subunit beta-1.

